# Effects of thermal stress on amount, composition, and antibacterial properties of coral mucus

**DOI:** 10.1101/368332

**Authors:** Rachel M. Wright, Marie E. Strader, Heather M. Genuise, Mikhail V. Matz

## Abstract

The surface mucus layer of reef-building corals supports several essential functions including feeding, sediment clearing, and protection from pathogenic invaders. For the reef ecosystem, coral mucus provides energy to support heterotrophic benthic communities. Mucus production represents a substantial metabolic investment on behalf of the coral: as much as half of the fixed carbon supplied by the corals’ algal symbionts is incorporated into expelled mucus. In this study, we examined if bleaching (disruption of the coral–algal symbiosis) has the potential to indirectly disturb reef ecosystem function by impacting the nutritional composition of coral mucus. In a controlled laboratory thermal stress challenge, visibly paled corals produced mucus with higher protein and lipid content and increased antibacterial activity relative to healthy corals. These results are likely explained by the expelled symbionts in the mucus of bleached individuals. This study illuminates how the immediate effects of coral bleaching could impact the reef-ecosystem indirectly through modulation of available nutrients within the ecosystem.

## Introduction

Coral bleaching results from the breakdown of the symbiosis between a coral host and its algal symbiont, *Symbiodinium*. Rising sea surface temperatures have increased the global risk of coral bleaching to alarming levels (Hughes et al. 2018), highlighting the need to understand the impacts of bleaching on both coral populations and the ecosystems they support. The direct impacts of bleaching on the animal host and algal symbiont are well studied. For example, coral bleaching has been shown to down-regulate genes related to host immunity (Pinzon et al. 2015) and alter host metabolism (Kenkel, Meyer, and Matz 2013; Rodrigues and Grottoli 2007). Symbionts expelled during bleaching produce elevated amounts of reactive oxygen species, but are otherwise physiologically similar to endosymbionts (Nielsen, Petrou, and Gates 2018). Numerous studies have also examined the indirect effects of coral bleaching and consequent mortality on community structure and function. For example, mass coral bleaching induces shifts in reef-fish assemblage structure and alters recruitment success (Richardson et al. 2018; Booth and Beretta 2002). However, the impact of coral bleaching on the nutrient cycle involving coral mucus is largely unknown.

Coral mucus is a complex mixture of proteins, lipids, and carbohydrates that is produced by mucocytes in the coral epidermal layer and secreted by coral surface tissues. Up to about half of the photosynthetically fixed carbon supplied by a coral’s algal symbiont is expelled as mucus (Crossland 1987; Crossland, Barnes, and Borowitzka 1980; Davies 1984). This coral surface mucus layer acts as a defense against desiccation and pathogens for the coral (reviewed in (Brown and Bythell 2005), and is also released into the water column where it traps suspended particles and acts as an energy source for benthic communities (Wild et al. 2004). Given the integral role of photosynthetically fixed carbon in producing coral mucus, it is predicted that coral bleaching events will reduce mucus production and subsequently impact the flow of energy throughout the reef ecosystem (Bythell and Wild 2011). However, the extent to which coral bleaching shifts the nutritional composition and function of coral mucus is yet to be characterized.

In the Florida Keys, annual mass bleaching events are predicted to begin by the mid-century (Manzello 2015). Currently, multiple anthropogenic factors including thermal stress, increased storms, and disease outbreaks have led to a near 80% decline in reef cover in the Florida Keys since the 1980s (Williams and Miller 2011; Williams, Miller, and Kramer 2008; Porter et al. 2001). In particular, corals in the genus *Acropora* have faced some of the most dramatic declines in this region. The staghorn coral, *Acropora cervicornis*, has been selected as a focal species for multiple active restoration programs, such as the Coral Restoration Foundation, due to its relatively fast asexual growth through fragmentation. As restoration efforts aim to replenish stands of *A. cervicornis*, it is critical to assess this species’ greater role in coral reef ecosystem. This study aims to characterize how coral mucus changes during acute thermal stress in *A. cervicornis* and to determine the potential consequences of this change on the coral reef ecosystem.

## Materials & Methods

### Corals

Fifteen *Acropora cervicornis* genets (n = 3 fragments per genet) were shipped from the Coral Restoration Foundation (Key Largo, Florida USA, Project ID CRF-2016–021) on 7 September 2016 to the University of Texas at Austin. Upon arrival, corals were immediately tagged with colored zip ties to uniquely identify each genet and allowed to recover for 12 days in artificial seawater (ASW; 30–31 ppt) at 25°C under 12000K LED lights on a 12h/12h day/night cycle. Corals were fed weekly with Ultimate Coral Food (Coral Frenzy, LLC).

### Experimental conditions

Coral fragments were partitioned into experimental and control tanks. One genet (U10) experienced mortality during the recovery period, so only one U10 fragment remained when the experiment began. For all other genets, one fragment was placed in a control tank (26°C) and one fragment was placed in an experimental tank. Any remaining fragments from the shipment of n = 3 per genet were retained in a holding tank, though many genets developed tissue loss or experienced damage on a single fragment during shipping. The single remaining U10 fragment was placed in the experimental tank. The temperature in the experimental tank was ramped from 26°C to 31°C over 33 hours. High summer temperatures in the Florida Keys often reach 31°C (Manzello 2015). Therefore, a 31°C heat treatment was chosen to represent an ecologically relevant stressor.

After corals had been exposed to experimental conditions for four days, corals appeared visibly pale relative to initial photographs and paired control fragments (Figure 1, Figure 3A). At this time, the temperature in the experimental tank was reduced to 26°C over 6 hours.

**Figure 1:**
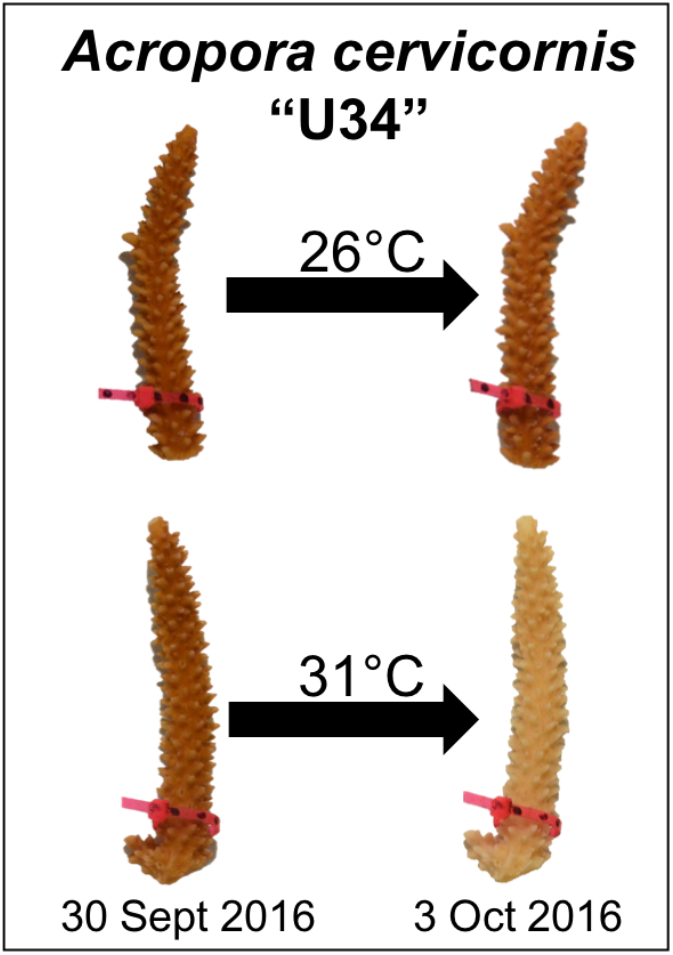
Experimental design and representative coral image. Corals were maintained in either control (26°C) or experimental (31°C) conditions for four days. Paling was observed for fragments in the experimental treatment, but not under control conditions.

### Image analysis

Prior to the experiment, photographs of each fragment were taken using a Nikon D5100 camera. Images of the front and back of each fragment were taken using the same camera, settings, and lighting each day of the experiment. Brightness values in images were measured for the front and back sides of each fragment using image analysis software (ImageJ, (Schneider, Rasband, and Eliceiri 2012). Corals become brighter (paler) as their symbioses with pigmented *Symbiodinium* break down. Therefore, changes in coral brightness reflect changes in *Symbiodinium* densities (Winters et al. 2009). A standard curve of brightness values was constructed using standard Coral Health Charts that were included in each image. Brightness values were standardized to color cards to normalize for any minor differences in lighting across days.

### Mucus collection

After the experiment, each coral fragment was placed within a pre-weighed 50 mL conical tube containing 5 mL ASW from the respective tank. Tubes were placed on their sides and secured to a gently rocking incubator plate (135 RPM, 28°C) for 20 minutes, rotating the tubes every 5 minutes to ensure that all sides of the coral fragment were submerged in water. After rocking, fragments were inverted dry above the liquid in the conical for 20 minutes and lightly centrifuged (200 RPM) for 2 minutes to pull down mucus adhering to the surface of the coral, modified from (Wild et al. 2004). The volume and weight of mucus from each fragment was measured and stored at –80°C. Coral fragments were returned to their tanks. The mucus collection procedure was repeated six days later, exactly as described above. During mucus collection, algal cells were clearly visible in some samples. All mucus aliquots were briefly centrifuged to remove the algal pellet before the experiments described below.

### Mucus composition

Total protein was measured following the Coomassie (Bradford) Protein Assay Kit (Thermo Scientific, Waltham, MA, USA). Total carbohydrate was measured using the Total Carbohydrate Quantification Assay Kit (Abcam, Cambridge, UK). Total lipids were extracted and the dry weights of each mucus sample were measured. A standard curve was prepared using reagent grade cholesterol in a 2:1 chloroform:methanol mixture and an aliquot of 2:1 chloroform:methanol was added to each sample tube. After mixing, the solvent was evaporated from all standard and sample tubes on a heat block at 90°C. Concentrated sulfuric acid was added to each tube, then incubated at 90°C for 20 minutes. Samples were cooled, then plated in triplicate into wells of a 96-well plate. Background absorbance was measured at 540 nm. After incubating each sample with 50 uL of vanillin-phosphoric acid for 10 minutes, absorbance was measured again at 540 nm. The concentrations of protein, carbohydrate, and lipid in the mucus, estimated using standard curves, were normalized to the volume of mucus expelled and the surface area of the fragment.

### Antibacterial activity

Cultures of laboratory *E. coli* (K-12) were grown overnight in LB, then washed twice in sterile ASW to remove remaining culture media. Coral mucus (140 µL) and washed *E. coli* culture (60 uL) was added to triplicate wells in 96-well plates. The covered plates were incubated at 37°C for 12 hours. Every 30 minutes the plate was shaken and the absorbance at 600 nm was measured.

### Coral surface area

Fragment surface area was estimated using a 3D scanner and accompanying ScanStudioPro software (NextEngine, Santa Monica, CA, USA). Each scan was completed using a 360 degree scan with 16 divisions and 10,000 points/inch^2^. Scans were then trimmed, polished to fill holes, fused and then surface area was estimated based on a size standard.

### Real-time quantitative PCR

The forward primer 5’-TCTGTACGCCAACACTGTGCTT-3’ and reverse primer 5’-AGTGATGCCAAGATGGAGCCT-3’ was used to amplify the *Acropora cervicornis* actin sequence as developed in (Winter 2017). The forward primer 5’-GTGAATTGCAGAACTCCGTG-3’ and reverse primer 5’-CCTCCGCTTACTTATATGCTT-3’ was used to amplify the *Symbiodinium* ITS2 sequence. Primer pair specificity was verified by gel electrophoresis and melt curve analysis of the amplification product obtained with *A. cervicornis* holobiont DNA. Primer efficiencies were determined by amplifying a series of four-fold dilutions of *A. cervicornis* holobiont DNA and analyzing the results using *PrimEff* function in the *MCMC.qpcr* package (Matz, Wright, and Scott 2013) in R. Briefly, C_T_ (threshold cycle) results were plotted as CT vs. log_2_[DNA], and amplification efficiencies (amplification factor per cycle) of each primer pair were derived from the slope of the regression using formula: efficiency = 2^-(1/slope)^ (Pfaffl 2001).

Mucus aliquots were centrifuged to remove any cell debris. A 14 µL aliquot of coral mucus was combined with SYBR Green PCR Master Mix (Applied Biosystems), 1.5 µM forward and reverse primers, and water. The Roche LightCycler 480 system was used to carry out the PCR protocol (95°C for 40 seconds, then 40 cycles of 60°C for 1 minute and 72°C for 1 minute) and detect the fluorescence signal.

### Statistics

All statistical analyses were performed in R (3.4.0, (R Core Team 2017)). The MCMCglmm package (Hadfield 2010) was used to explain variation in coral color and mucus composition, with treatment as a fixed effect and genotype as a random effect. The nlme package (Pinheiro et al. 2017) was used for time-series analysis of antibacterial activity, with time, treatment, and their interaction as fixed effects and plate well as a random effect. Experimental data and R scripts are included as Supplemental Data S1 and S2, respectively. Data and scripts can also be accessed on GitHub: https://github.com/rachelwright8/bleached_coral_mucus.

## Results

### Coral bleaching and mucus collection

After four days in the experimental treatment at 31°C, corals paled significantly compared to corals in the control condition (β = −1.16, p < 0.001, Figure 3A). Corals from both treatments produced similar amounts of mucus by volume (β = 0.01, *p* = 0.25, Figure 2A) and weight (β = 0.14, *p* = 0.27, Figure 2B). After a six-day recovery period, corals in both treatments produced significantly less mucus by volume (β = −0.14, *p* < 0.001, Figure 2A) and weight (β = −1.1, *p* < 0.001, Figure 2B) compared to the first time point.

**Figure 2:**
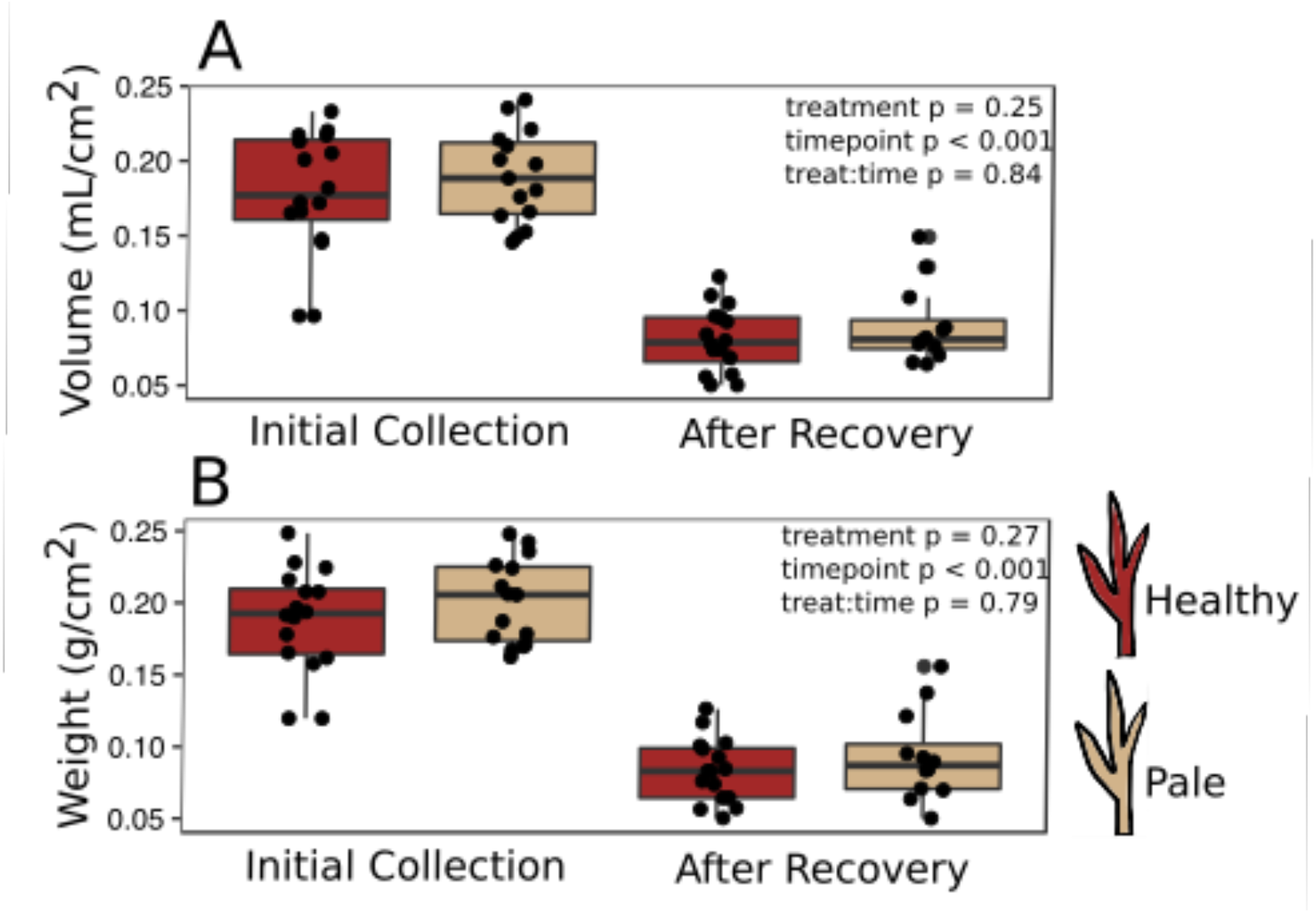
Mucus production. Mucus was collected immediately after paling was observed (“Initial”) and six days after the challenged corals were returned to control conditions (“Recovery”). The volume (**A**) and weight (**B**) of the recovered mucus was normalized to the surface area of the coral fragment.

### Mucus biochemistry

Although heat-stressed corals were visibly pale, the mucus produced by these fragments contained significantly more total protein (β = 2.1, *p* < 0.001, Figure 3B) and total lipid (β = 15.7, *p* = 0.02, Figure 3C). There was also a marginally significant increase in carbohydrate content in mucus from pale corals (β = 0.64, *p* = 0.10, Figure 3D).

**Figure 3:**
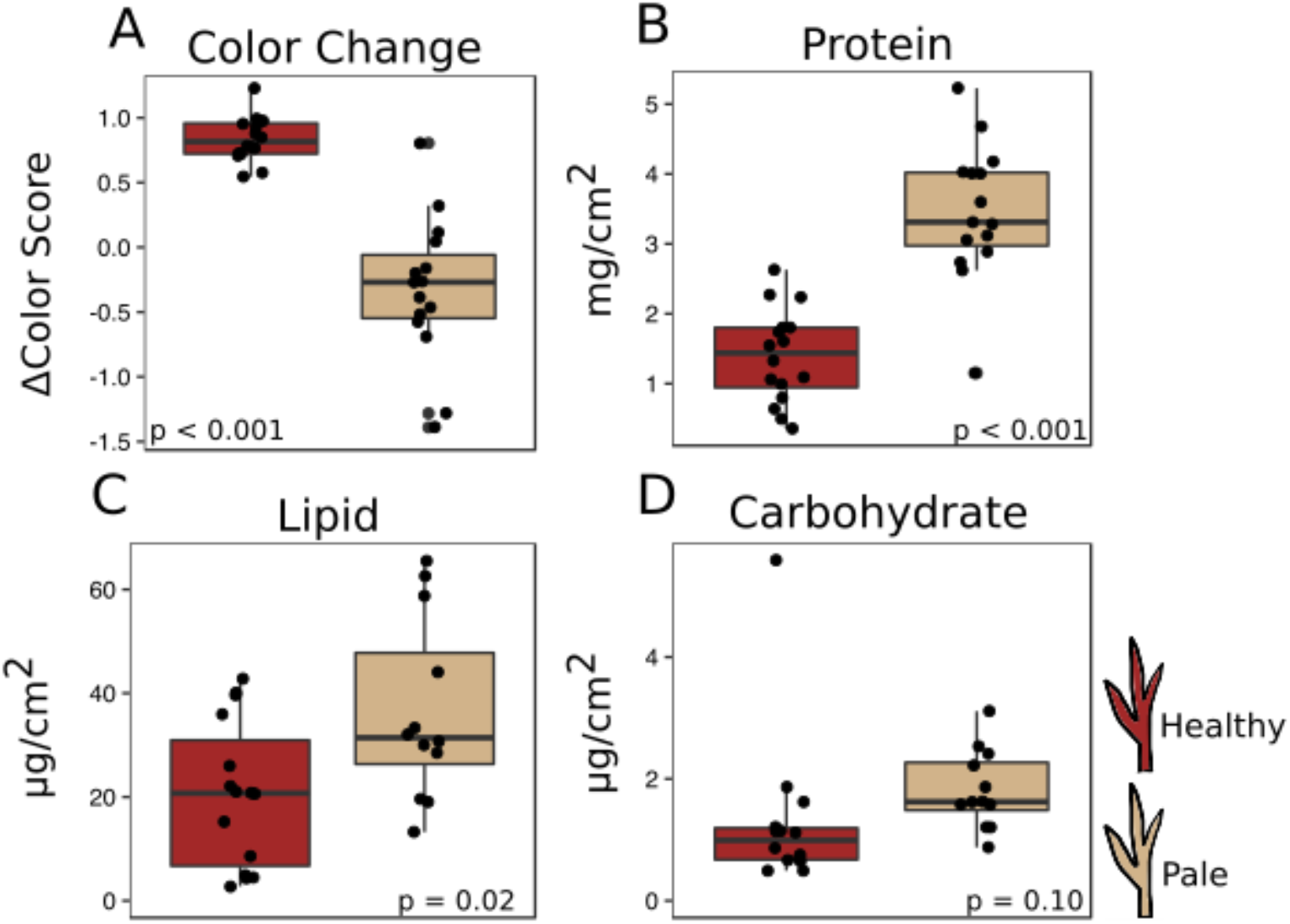
Effects of heat stress. Dark red boxes represent fragments in control conditions, beige boxes represent heat-stressed fragments. **A**: Effect on coral color (decrease in color score indicates bleaching). (**B–D**): Effects on mucus composition: protein (**B**), in mg/cm^2^ fragment surface area, and lipid (**C**) and carbohydrate (**D**), in µg/cm^2^ fragment surface area.

### Mucus antibacterial activity

Antibacterial activity increased in thermally stressed corals from the experimental treatment relative to healthy corals (ANOVA *p* = 0.0067, Figure 4). Absorbance at 600 nm, which reflects bacterial density, decreased throughout the incubation period in all mucus samples. However, bacterial density declined significantly faster in mucus samples from heat-stressed corals.

**Figure 4:**
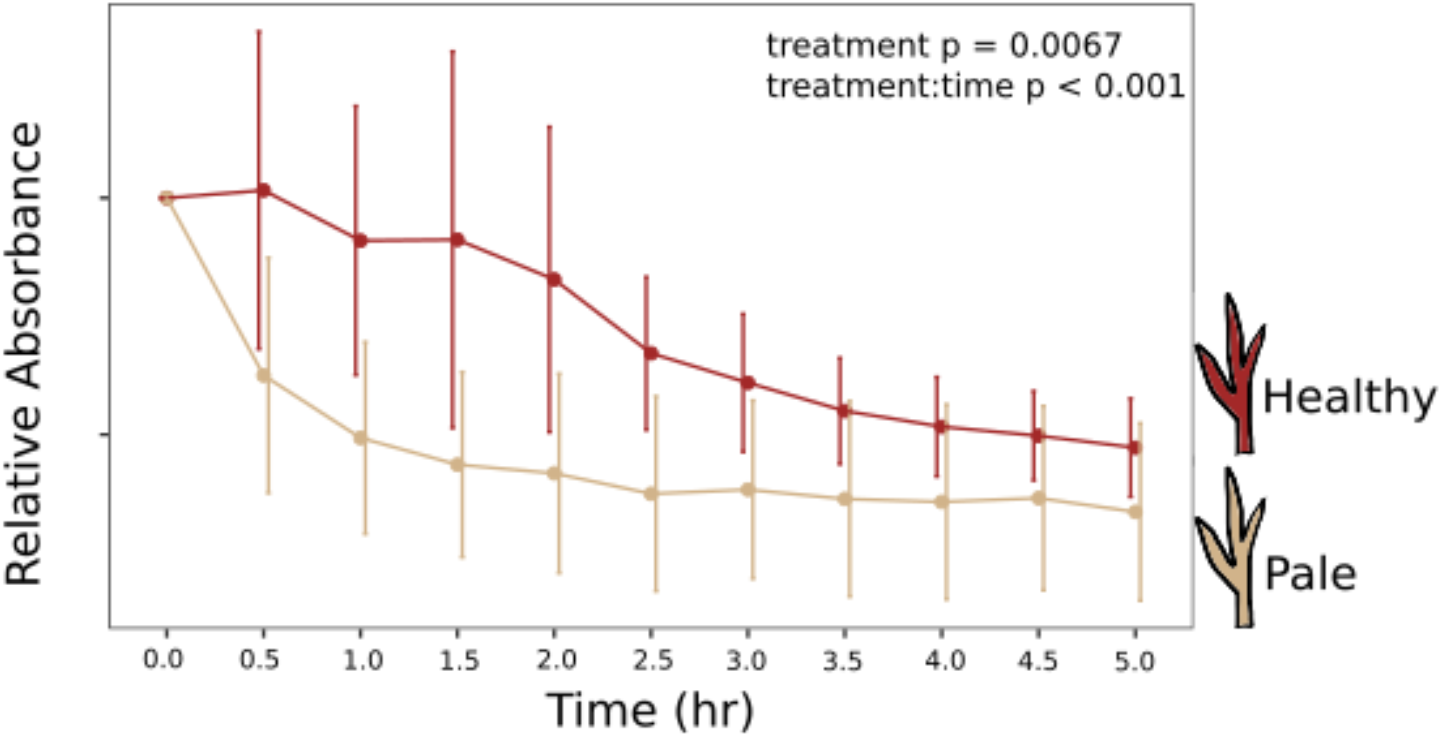
Antibacterial activity of coral mucus. Absorbance at 600 nm reflects the density of inoculated bacteria in coral mucus samples.

### Presence of host and symbiont DNA in mucus

Real-time quantitative PCR (qPCR) was performed to determine the relative abundances of coral- and *Symbiodinium*-derived DNA sequences present in the mucus released by healthy and heat-stressed corals. Primers were designed to target a coral-specific actin gene and the *Symbiodinium* ITS2 region. Presumably, copies of the coral-specific actin gene would represent lysed coral cells, while copies of the ITS2 region would represent material released from *Symbiodinium* cells in the mucus.

Few coral-specific actin copies were detected and no ITS2 sequences were present in the mucus of unchallenged, healthy corals (Figure 5). However, mucus released by heat-stressed, pale coral fragments contained abundant copies of both coral- and symbiont-derived sequences (Figure 5).

**Figure 5:**
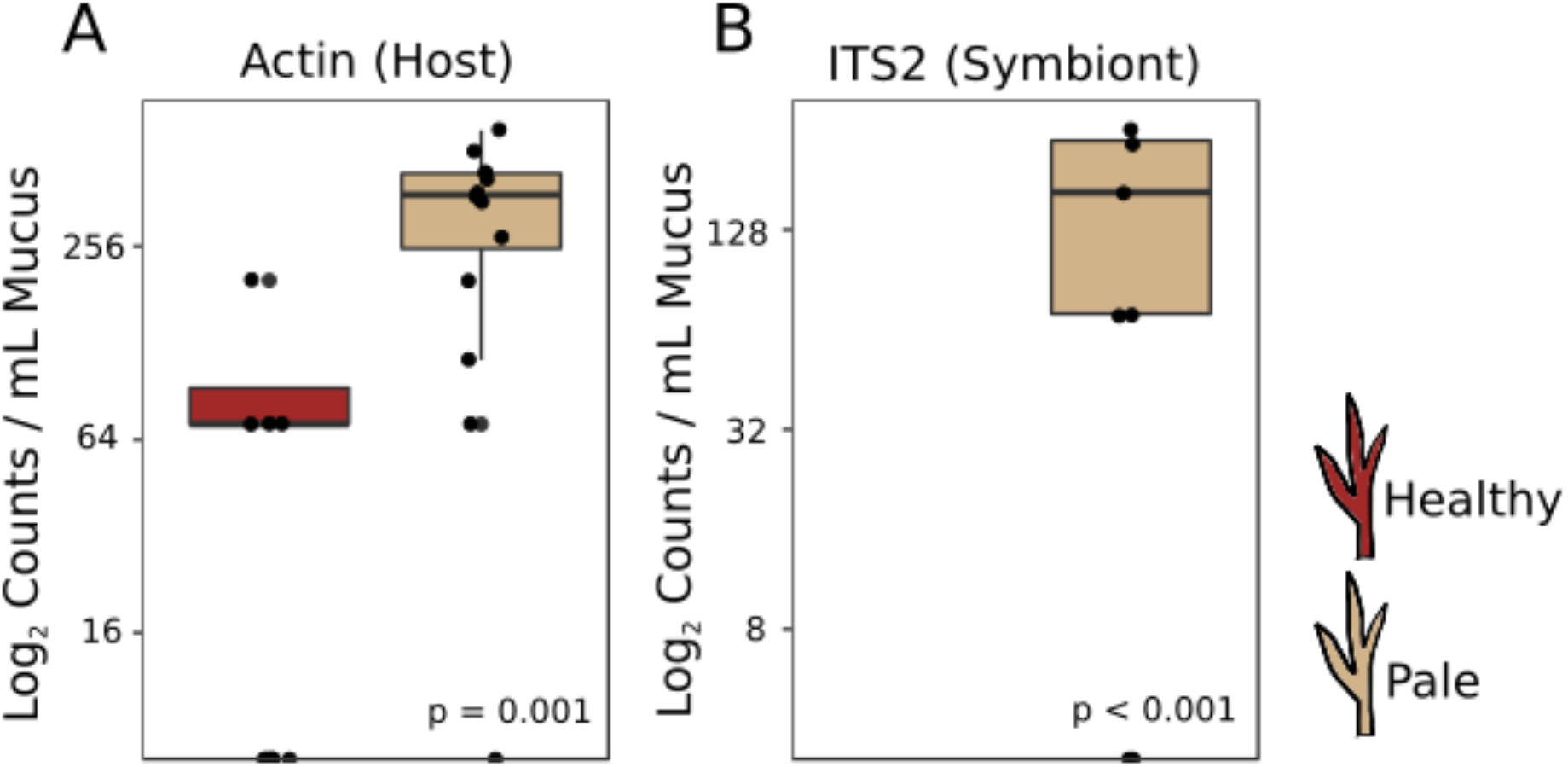
*Symbiodinium* and coral DNA in mucus. Real-time quantitative PCR detected copies of *A. cervicornis* actin (**A**) or *Symbiodinium* ITS2 (**B**) in the coral mucus of healthy and pale corals.

## Discussion

### Coral mucus stores take a long time to replenish

Mucus volumes released during the initial collection (0.18±0.04 mL/cm^2^, or about 1.8 L/m^2^) are consistent with daily mucus release values previously reported for submerged acroporids (1.7 L/m^2^ in Wild et al. 2004). However, following a six-day recovery period, less than half of the original mucus volume was collected (46.8±13.2%, *p* < 0.001, Figure 2A), suggesting that mucus stores were not completely replenished in this amount of time. The methods used in this study to extract mucus left the coral nubbins completely dry. Therefore, the results presented here represent the total mucus attached to a coral at any one time rather than the amount of mucus may be naturally released into the water column daily. These results emphasize the importance of measuring mucus release over time to confidently estimate daily release rates and predict daily energetic flow throughout the reef ecosystem.

### Stressed corals produce mucus high in protein and lipid

Given the energetic cost of producing mucus (Riegl and Branch 1995), it is reasonable to predict that corals with low densities of autotrophic symbionts would produce less, or lower nutritional quality, mucus than healthy corals. We found no difference in the quantity of mucus produced by stressed corals compared to healthy corals after four days of heat stress or after a six-day recovery period (Figure 2), suggesting that the quantity of mucus produced a coral is relatively unaffected by thermal stress and that mucus stores cannot be replenished within a week. Surprisingly, we found higher protein and lipid content in mucus from pale corals relative to mucus produced by healthy corals (Figure 3). *Symbiodinium* store reserve energy as lipid droplets and starch granules that are translocated from the algal membrane to coral cells in a healthy coral–algae symbiotic relationship (Patton and Burris 1983). We observed a pellet of *Symbiodinium* cells in the mucus of thermally stressed corals, but not in mucus produced by healthy corals. Though *Symbiodinium* cells were pelleted and removed from all mucus collections, extracellular lipid droplets would remain in the mucus and represent a potential explanation for the increased abundance of lipids in mucus from stressed corals. Likewise, proteins and lipids released from damaged *Symbiodinium* and host cells would also be present in the mucus of stressed corals. This particular possibility is supported by finding of both coral and *Symbiodinium* DNA in the mucus of stressed corals (Figure 5). Future studies should measure long-term effects of bleaching to determine the duration of this observed enrichment in coral mucus quality following thermal stress.

### Stressed corals produce mucus with high antibacterial activity

Surprisingly, this study found that mucus collected from stressed coral fragments eliminated bacteria faster than mucus from healthy fragments from matched genotypes (Figure 4). Antibacterial activity of coral mucus is attributed to antimicrobial substances produced by commensal microbes living on the coral surface (Nissimov, Rosenberg, and Munn 2009; Shnit-Orland and Kushmaro 2009). In contrast to our results, a previous study found reduced antibacterial activity in mucus collected from *A. palmata* during a summer bleaching event in the Florida Keys (Ritchie 2006). The discrepancy in findings could be attributed to the timing of collections. Mucus in this study was collected as soon as corals became pale, whereas the 2005 study collected mucus after corals in the Florida Keys had been experiencing high levels of thermal stress and bleaching for about a month (Eakin et al. 2010). Long-term thermal stress is known to promote coral disease by altering bacterial pathogenicity and host susceptibility (Bruno et al. 2007; Maynard et al. 2015). In our short-term bleaching conditions, the increased protein and lipid content in the mucus (Figure 3B-C) may have temporarily improved the antibacterial activity of commensal microbes that exist in the coral mucus. Another possibility is that the expelled *Symbiodinium* themselves released some antimicrobial products. Though the mechanism is unclear, *Symbiodinium* do appear to play a role in a coral’s ability to manage immune stress and regulate microbial communities (Littman, Bourne, and Willis 2010; Rouzé et al. 2016; Wright et al. 2017).

## Conclusions

Our results show that thermal stress does not significantly affect the volume of mucus produced by *A. cervicornis* immediately following a bleaching event. Surprisingly, stressed corals produced mucus with higher protein content, higher lipid content, and increased antibacterial activity relative to unstressed controls. Additional lipids and proteins likely come from *Symbiodinium* and host cells damaged during bleaching rather than from additional investment by the coral host. Elevated nutritional value of mucus released from bleaching corals could have significant consequences for the reef’s nutrient cycle, while changes in both nutritional composition and antibacterial properties of the mucus should strongly affect coral-associated microbes and, as a consequence, coral disease susceptibility. Future experiments should investigate longer-term effects of thermal stress on mucus production and content to further investigate reef-wide consequences of coral bleaching.

## Acknowledgements

We thank the Coral Reef Foundation for providing coral specimen and Sarah Davies for measuring coral surface areas.

## Supplemental Files

Data S1: Excel file containing experimental data.

Data S2: R script for analyzing data.

